# Improving characterization of understudied human microbiomes using targeted phylogenetics

**DOI:** 10.1101/2019.12.16.879023

**Authors:** Bruce A Rosa, Kathie Mihindukulasuriya, Kymberlie Hallsworth-Pepin, Aye Wollam, John Martin, Caroline Snowden, Wm. Michael Dunne, George Weinstock, Carey-Ann D. Burnham, Makedonka Mitreva

**Author notes:** Correspondence should be addressed to Makedonka Mitreva. Bruce A Rosa –, Kathie Mihindukulasuriya –, Kymberlie Hallsworth-Pepin –, Aye Wollam –, John Martin –, Caroline Snowden –, Wm. Michael Dunne, Jr. –, George M. Weinstock –, Carey-Ann D. Burnham –, Makedonka Mitreva –.

## Abstract

Whole genome bacterial sequences are required to better understand microbial functions, niches-pecific bacterial metabolism, and disease states. Although genomic sequences are available for many of the human-associated bacteria from commonly tested body habitats (e.g. stool), as few as 13% of bacterial-derived reads from other sites such as the skin map to known bacterial genomes. To facilitate a better characterization of metagenomic shotgun reads from under-represented body sites, we collected over 10,000 bacterial isolates originating from 14 human body habitats, identified novel taxonomic groups based on full length 16S rRNA sequences, clustered the sequences to ensure that no individual taxonomic group was over-selected for sequencing, prioritized bacteria from under-represented body sites (such as skin, respiratory and urinary tract), and sequenced and assembled genomes for 665 new bacterial strains. Here we show that addition of these genomes improved read mapping rates of HMP metagenomic samples by nearly 30% for the previously under-represented phylum *Fusobacteria*, and 27.5% of the novel genomes generated here had high representation in at least one of the tested HMP samples, compared to 12.5% of the sequences in the public databases, indicating an enrichment of useful novel genomic sequences resulting from the prioritization procedure. As our understanding of the human microbiome continues to improve and to enter the realm of therapy developments, targeted approaches such as this to improve genomic databases will increase in importance from both an academic and clinical perspective.

**Importance:** The human microbiome plays a critically important role in health and disease, but current understanding of the mechanisms underlying the interactions between the varying microbiome and the different host environments is lacking. Having access to a database of fully sequenced bacterial genomes provides invaluable insights into microbial functions, but currently sequenced genomes for the human microbiome have largely come from a limited number of body sites (primarily stool), while other sites such as the skin, respiratory tract and urinary tracts are under-represented, resulting in as little as 13% of bacterial-derived reads mapping to known bacterial genomes. Here, we sequenced and assembled 665 new bacterial genomes, prioritized from a larger database to select under-represented body sites and bacterial taxa in the existing databases. As a result, we substantially improve mapping rates for samples from the Human Microbiome Project and provide an important contribution to human bacterial genomic databases for future studies.

## Introduction

As sequencing technology improves, the increased practicality of genomic analysis has allowed for new inquiries into the workings of the human microbiome. Already, research has characterized the microbial communities of diverse body sites and elucidated the role that microbiota may play in conditions including type 2 diabetes, cardiovascular disease, obesity, cancer, and autism (1–4). Future research aspires to manipulate microbial communities as a prophylactic or therapeutic approach to prevent and/or control various infectious and disease states.

The data for these microbiome studies generally take the form of either 16S ribosomal RNA gene sequences or metagenomic shotgun sequencing. The former is frequently used for its efficiency and affordability, requiring only marker genes to be known in order to align and annotate reads. However, exclusive use of this type of data has become limiting to researchers who wish to understand a more complete picture of microbial functions and metabolism within a niche. Metagenomic shotgun sequencing can provide answers to these kinds of question, but it relies upon the use of reference genome for either alignment or k-mer comparison of genomes intermixed from hundreds to thousands of different taxa (5).

In the past, there have been several efforts to compile a database of microbial genomes. Most notably, the MetaHIT (6) and Human Microbiome Project (HMP) (7) gene catalogs made available catalogs based upon over a hundred European and American samples, respectively. In 2014, Li et al amassed this data in addition to data from other international studies to create an integrated gene catalog for the human gut microbiome (IGC) (8). This catalog contained almost ten million non-redundant genes, but it still remains far from comprehensive. Most of these species were cultured from healthy patients whose microbial communities likely differ from those of disease states (9). Additionally, a disproportionate amount of the available reference genomes belong to gut and urogenital microbiota, neglecting other taxa that might be pertinent to studies of skin, vaginal, or respiratory microbiomes (10). In fact, a 2014 study examining the effect of genomic database composition on metagenomic read mapping found that with existing databases, read mapping varied significantly between body sites. The highest-performing areas (posterior fornix) mapped as many as 92% of reads, but samples in the lowest-performing areas (skin) mapped as low as 13% (11). Finally, microbiome composition varies not only between sites on an individual, but also between individuals in diverse environments. Studies have demonstrated significant differences in the microbiome composition of individuals from different geographic areas, suggesting that a broader spectrum of reference genomes may be necessary for applicability of microbiome analytic techniques on a global scale (12).

As studies continue to unearth the variety of microbiomes found in multiple countries or a single individual, as well as the importance of tracing new bacterial strains during a public health outbreak, it becomes increasingly important to populate the bacterial phylogeny while representing diverse body habitats by generating high-quality reference genomes to publicly available databases. In this study the sequencing and analyzes of 10,000 full-length 16S rRNA gene sequences identified underrepresented phylogenetic lineages that were sequenced at a whole genome level. Analyses of the 665 genomes illuminated the importance of taxonomic prioritization in new genome discovery and demonstrated the effects of unbiased database on microbiome characterization. The approach and the resources are of importance at both the academic and clinical research level.

## Results and Discussion

### Composition of sequenced genomes

A total of 4,546 full-length 16S rRNA gene sequences were interrogated based on: (i) their 16S classification, prioritizing novel or under-represented taxonomic groups in existing public databases; (ii) their body site source, prioritizing under-represented body sites (such as blood, peritoneal fluid, and wounds); (iii) the quality and availability of the source DNA. The analysis resulted in 665 strains to be selected for whole genome sequencing, assembly and annotation represented 15% of the candidate samples initially screened for novelty.

Before and after prioritization, samples were taken from over 15 different body sites, with the largest taxonomic representation drawn from skin, respiratory, and urinary tract cultures (Figure 1A). Prioritization caused a relative increase of at least 300% in sparsely sampled areas such as joint and bone, while relative representation of samples from medical hardware, urinary tract and unknown locations were reduced, perhaps due to the homogeneity of bacterial communities that has previously been observed (13, 14). The gut microbiota was under-represented among pre-prioritized and prioritized samples, since it is the most well characterized of the body locations (10). Additionally, the prioritization considered phylogenetic clustering, with the intention to avoid over-selecting the same strains of bacteria, regardless of species or sample site origin (99% identity over 95% length). This resulted in a wider array of unique taxa being sequenced than would have been selected without considering phylogeny.

**Figure 1:**
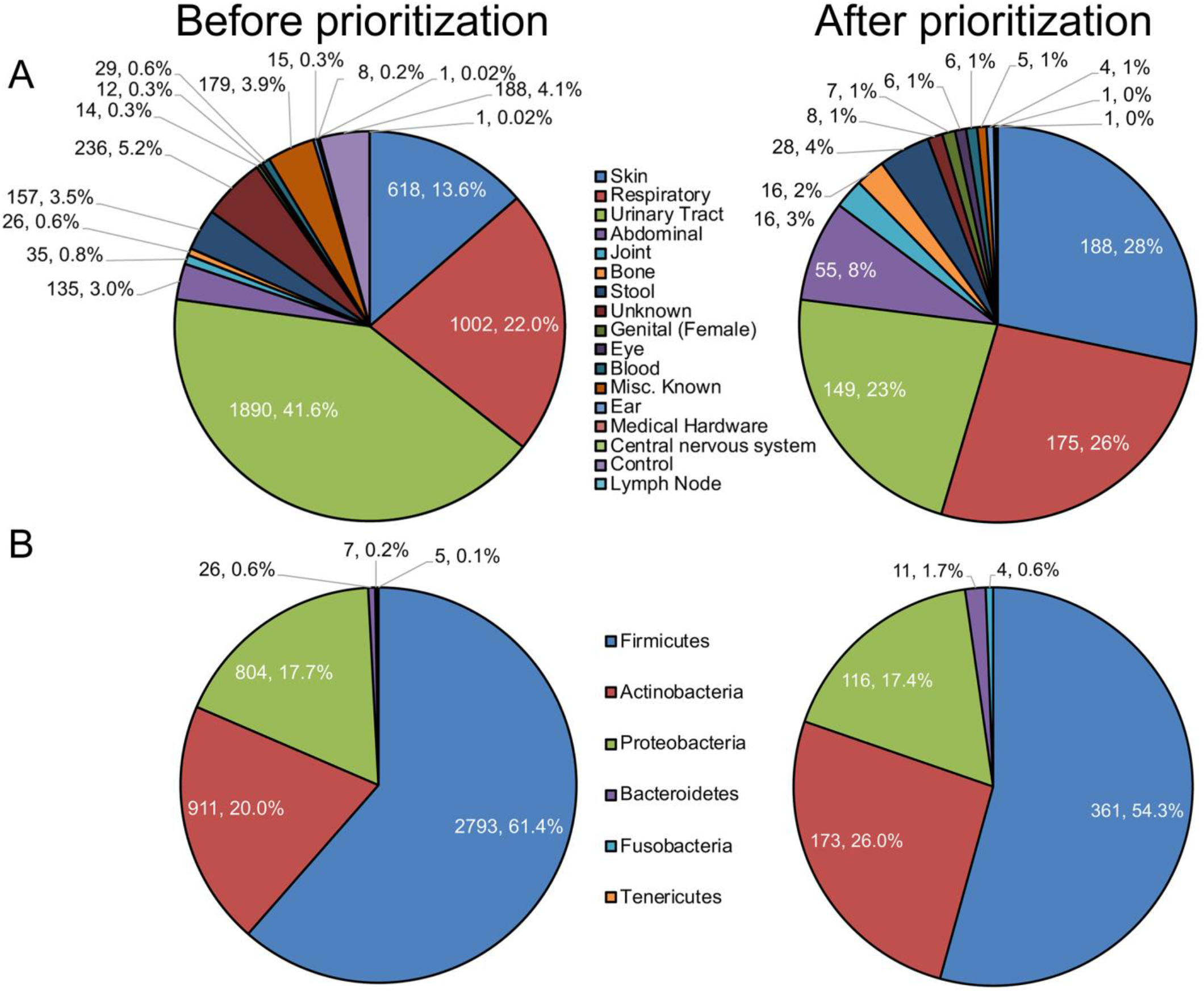
Composition of isolates before prioritization (left) and after (right). **A**) Isolate categorization by body habitat. **B**) Isolate categorization by phylogeny at a phyla level.

The majority of samples before and after prioritization were part of the *Firmicutes* phylum, followed in abundance by the *Actinobacteria* and *Proteobacteria* phyla. *Bacteroidetes, Fusobacteria* and *Tenericutes* were sparsely represented in samples (Figure 1B). After prioritization, relatively fewer *Firmicutes* samples were chosen for sequencing, since studies historically have identified primarily taxa in the *Bacteroidetes* and *Firmicutes* phyla, which tend to be more abundant in the gut (10). Instead, the relative representation of *Actinobacteria* in samples to be sequenced increased after prioritization. *Actinobacteria* is an important component of several different body location microbiomes, including skin folds (15) and the poorly-characterized eye mucosal microbiome (16). In addition to *Fusobacteria*, it also comprises a notable part of oral microbiomes such as that of the tongue dorsum or supragingival plaque (16). As the literature regarding these areas continues to expand, further characterization of their microbiomes will require genomic libraries including more taxa within the *Actinobacteria* or *Fusobacteria* phyla, which may not have appeared in earlier gut microbiome studies. This is becoming increasingly important as these taxa have been implicated in a number of health and disease states, such, as colon cancer (17–19).

Clustering of the prioritized isolates by 16S read counts showed clear assortment by phyla (Figure 2A). Some body sites tended to cluster within phyla; most notably, respiratory tract samples were primarily within the *Proteobacteria* phylum, while skin samples clustered within the *Firmicutes* phylum and *Bacilli* class (Figure 2B). It is interesting to note that these phyla are not those predominantly associated with the respiratory tract or skin, respectively. Studies suggest that the principal taxa in the respiratory tract lie within the *Bacteriodetes* and *Firmicutes* phyla (20), while skin samples vary greatly, but frequently contain *Actinobacteria, Proteobacteria* or bacteria within the *Staphylococcaceae* class of *Firmicutes* (15). The differences between our data clusters and the normal skin or respiratory tract communities reflect the efforts of our prioritization method to obtain sequences unique from the known biological landscape. These sequences may become particularly important in efforts to characterize disease-state deviations from the normal microbiome: for example, research suggests that asthma may be associated with enrichment of bacteria in the *Proteobacteria* phylum within which our prioritized respiratory samples tended to cluster (21). Characterization of such deviate taxa, as opposed to those that fall within the norm, may give us better resolution in our view of how physiological processes associate with changes in diverse microbiota.

**Figure 2:**
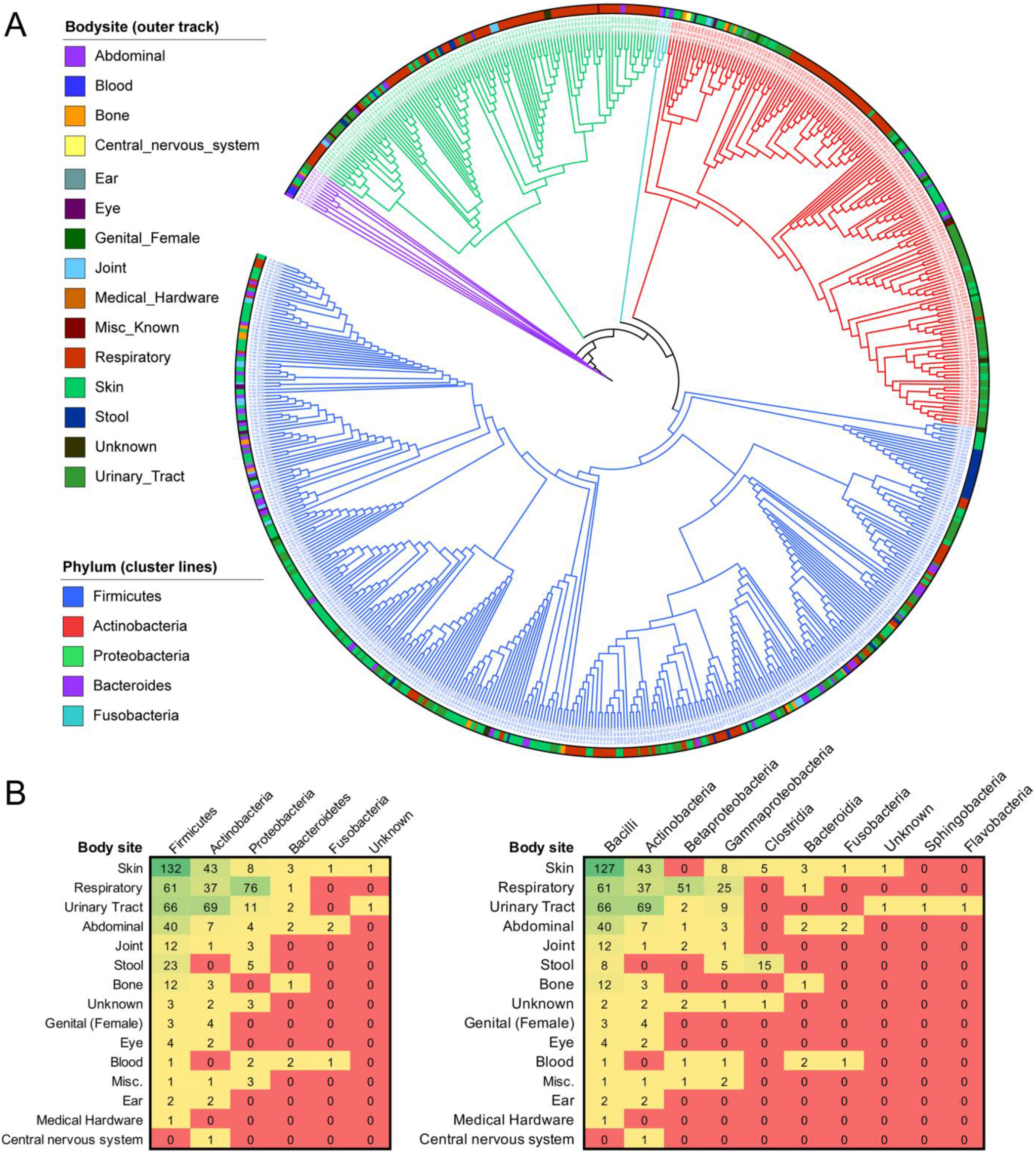
Grouping of the sequenced WUSC bacterial genomes based on body site, phyla, and class. **(A)** WUSC strains clustered based off of 16S read sequence similarity. Phyla is indicated by dendrogram branch color, while body habitat is indicated by the colored bars around the periphery of the image. **(B)** WUSC strains classified by body site and phylogeny at a phyla (left) or class (right) level. Counts indicate the number of strains in each category; color indicates the level of representation in each category, with green representing high counts and red representing low or none.

### Effect of database sequence augmentation on characterization of metagenomics shotgun sequences

Compared to publicly available genome databases alone (HMP + GenBank; 4,383 strains), the addition of our 665 novel genomes substantially increased genomic reads mapped for all 1,391 shotgun metagenomic samples from the HMP. By body site, the largest increases in reads mapped were observed for HMP samples from oral sites including the tongue dorsum and buccal mucosa (Figure 3A, Table 1). Besides GI samples, these sites were two of the most numerous found in the database, indicating a notable absolute as well as relative increase in amount of read mapping (Figure 3B). These sites were not the most frequent source of our novel sequences, but their read-mapping was increased significantly, indicating the importance of a multifactorial prioritization method taking into account the alignment and phylogenetic distinctness of candidate taxa in addition to their source.

**Figure 3:**
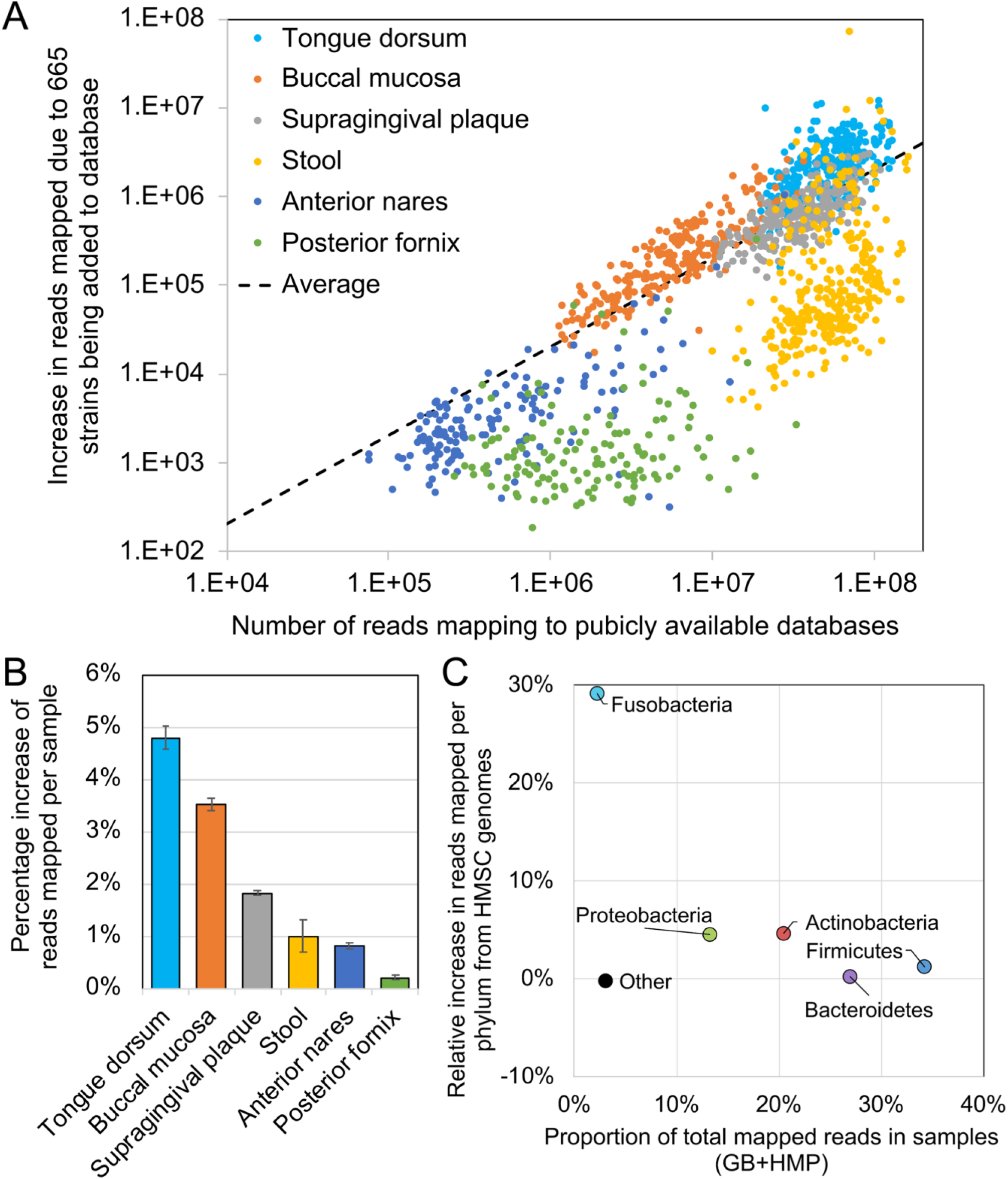
Increased characterization of metagenomic shotgun sequences by addition of genomes of novel phylogenetically distinct strains. **A**) Relative improvement of read mapping rates from HMP samples to the improved genome database. **B**) Number of total mapped reads per sample after novel reference genome sequence inclusion versus the absolute increase in mapped read after versus before novel sequence inclusion. Samples are colored by body site. The dotted line represents the average relationship between total number of mapped reads and absolute increase in mapped reads after novel reference genome sequence inclusion. **C**) The proportion of total mapped reads after novel sequence inclusion attributable to a given phyla versus the relative increase in reads mapped for that phyla after novel sequence inclusion. Phyla are represented as differently colored dots, with names given on the chart.

**Table 1:**
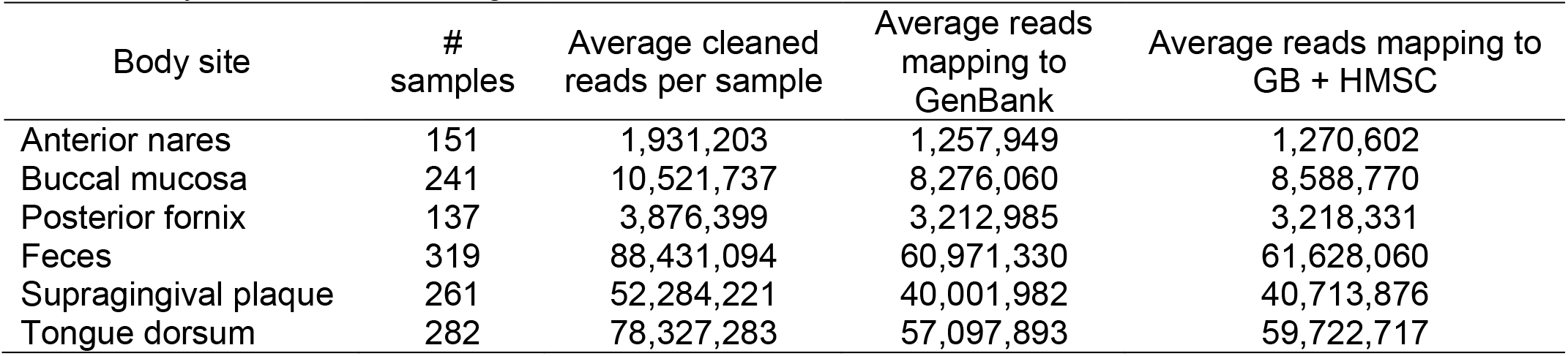
Average read mapping of samples grouped by body site to GenBank and novel HMSC genomes (GB + HMSC) versus GenBank alone. GenBank was considered the “baseline” database prior to this study’s addition of novel genome assemblies.

Taxonomically, the largest increase in reads mapped was observed for bacteria from the phylum *Fusobacteria*. This phylum has historically been poorly characterized in databases, so our sequence contributions increased mapped reads by almost 30% (Figure 3C). The phyla *Proteobacteria* and *Actinobacteria* showed modest increases in mapped reads, while the well-characterized *Firmicutes* and *Bacteroidetes* phyla showed only very small improvements. The improvements observed in read mapping for these taxa are likely related to the improvements we observed in read mapping for the oral mucosa, which contains greater proportions of *Fusobacteria, Actinobacteria* and *Proteobacteria* (22).

This overall improvement is unsurprising given the high representation of our novel genomes among diverse HMP samples. 183 of the 665 (27.5%) novel genomes sequenced had high representation in at least one of the HMP samples (≥50% breadth and ≥1X depth). In comparison, only 12.5% of the public database genomes had high representation (P<10^−5^ for enrichment of high-abundant strains; Figure 4).

**Figure 4:**
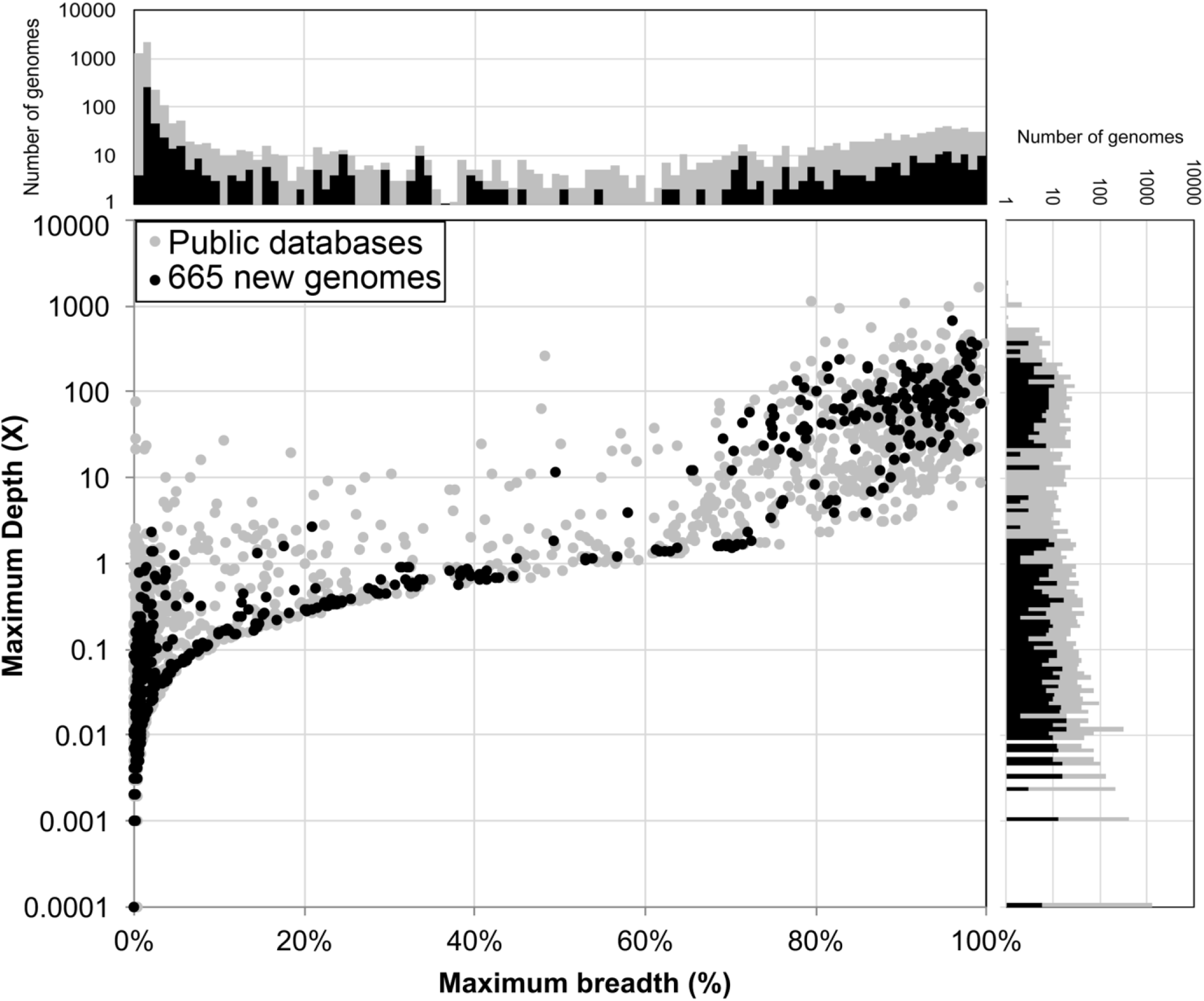
Novel genome assemblies (dark black) show increased representation in HMP samples, relative to sample size (>50% breadth, >1X depth, upper right quadrant of graph). Novel genomes are shown in black, pubic database genomes in dark gray. Numbers are not normalized for database size.

There are several possible reasons for the enrichment of our novel genomes in the HMP dataset relative to GenBank. First, our genomes were all sequenced from human samples, as opposed to many environmental samples that contribute to other public databases. For this reason, our results are particularly applicable to biomedical research endeavors. Second, our novelty by a phylogeny prioritization process, which selected a representative sequence from a cluster of similar sequences, emphasized variety in our rank list. Rather than simply sequencing all of the genomes that were determined to be the most novel, our process ensured sampling of novel genomes from many different taxa. Third, our selection of isolates from under-characterized body sites broadens the types of samples that would find representation in a database. At the moment, HMP statistics show a database composition primarily of GI and urogenital cultures, with several other body sites having fewer than ten reference genomes (23). For this reason, samples taken from body sites such as the eyes or respiratory system might contain genomes from our newly sequenced cohort but not from public databases.

If undertaken with focused sampling efforts, culturing and sequencing of isolates determined by our prioritization process could yield even greater improvement to known databases. Our isolates were taken from samples originally obtained from a variety of standard of care cultures types from patients suspected to have an infection, thus resulting in a variety of sample qualities. Poor quality or technical workability sometimes eliminated what would have otherwise been a highly-ranked isolate for sequencing. Additionally, our approach was still limited to those taxa that could be cultured in a laboratory setting using routine media commonly used in the clinical microbiology laboratory. There may be other taxa, especially among the variety of body sites in our study that are important but have heretofore not been sequenced due to their inability to grow in culture, or inability to grow with the culture techniques used as part of routine clinical microbiology.

Our results demonstrate the benefit of a targeted approach to the sequencing and assembly of new microbial genomes. It also contributed 665 novel genomes originating from diverse human habitats to public databases that can be utilized by other studies. If more endeavors similar to this one undertaken in the future, read mapping could be improved even more for microbial community samples spanning a variety of human habitats. As our understanding of the microbiome continues to improve and to enter the realm of therapy developments, such endeavors will increase in importance from both an academic and clinical perspective.

## Materials and Methods

### Sampling and culturing

Clinical specimens were submitted to the Barnes-Jewish Hospital Clinical Microbiology Laboratory and were processed using culture media, incubation atmosphere, and incubation time as described in the standard operating procedures for each specimen type. Bacterial isolates that morphologically resembled resident microbiota for the sample type cultured were selected, and a suspension of the microorganism was made for sequence based-analysis.

### Bacterial 16S rRNA genes were sequenced to identify novel bacterial strains associated with diverse habitats

The full-length of the 16S rRNA genes (16S) for each bacterial strain was obtained by sequencing three overlapping regions on the 3730 ABI platform. Primers used for the three amplicons were: V1-V3, 27F (AGAGTTTGATCCTGGCTCAG) and 534R (ATTACCGCGGCTGCTGG); V3-V5, 357F (CCTACGGGAGGCAGCAG) and 926R (CCGTCAATTCMTTTRAGT); and V6-V9, U968f (AACGCGAAGAACCTTAC) and 1492r-MP (TACGGYTACCTTGTTAYGACTT). A total of 10,787 microbial isolates with 16S reads generated on the Sanger 3730 platform were analyzed. Following analytical processing and removal of chimeric 16S reads (24), a phylogenetic step-wise approach was undertaken to prioritize the potential novel genomes to be sequenced. Comparison to publicly available 16S rRNA gene sequences was performed using a blast database that contained non-redundant 16S rRNA gene sequences from Silva v.115 (25) and RDP (26) training set 9. The 16S rRNA genes from the Washington University strain collection (WUSC) were blasted against this database, and any sequence with ≤ 97% identity and/or ≤ 90% coverage against any of the strains was considered potentially novel (225 total “novel” strains). Following this initial processing, of the 10,787 starting strains, 4,546 sequences (from 3,798 unique samples, due to the sequencing of technical replicates) met three criteria to advance to further analysis: successful 16S assembly (using the One Button Velvet assembly pipeline, Version 1.1.06 (27)), metadata completeness, and successful RDP classification(26).

To further prioritize isolates, we performed phylogenetic analysis to avoid repeatedly sampling very similar isolates, and to preferentially select novel sequences, such as those similar to known sequences, but isolated from a different body site. The 16S rRNA genes from the complete bacterial genomes from the HMP (whole genome shotgun-based sequencing; inclusive of the 4 HMP sequencing centers; McDonnell Genome Institute at Washington University School of Medicine, J. Craig Venter Institute, Baylor College of Medicine and the Broad Institute) and the GreenGenes GOLD database were clustered into a non-redundant database by clustering all sequences using 99% identity and 95% coverage (28), with the longest sequence per cluster used as a representative. The WUSC 16S rRNA genes from the 3,798 unique samples were then clustered with these representative sequences at 99% identity over 95% length. A total of 247 WUSC samples were determined to be contaminated due to the presence of multiple sequence replicates from the same sample being present in different clusters, leaving 3,551 samples for downstream prioritization.

The taxonomic classifications were used to split the sequences into taxonomic groups manageable for manual evaluation, by constructing phylogenetic trees for each taxonomic group using mothur (29). The 15 resulting phylogenetic trees were visualized using iTOL (30), and 16S rRNA sequences were considered to be novel by phylogeny if they did not cluster with the 16S rRNA gene of a sequenced bacterial genome or if they originated from different body sites than the sequenced isolate (262 samples). For simplicity in the prioritization, body site information was collapsed into a broader category when more detailed (e.g. abscesses from any body site were collapsed to “Abscess”).

The top available samples were prioritized for sequencing according to (i) all novel samples first (20 final samples sequenced); (ii) the top-ranked sample within each novel by phylogeny cluster (88); (iii) known cluster top representatives (51); (iv) novel by phylogeny samples from unique isolation sources, but which were not the top-ranked in their cluster (Only up to 5 per cluster, since there are some very large clusters; 128), and (v) novel by phylogeny samples, sorted by within-cluster rank, selecting for samples from rare or desirable body sites (355); (vi) known, unique isolation sources (23).

Overall, from the 10,787 starting 16S sequences in the WUSC database, 4,546 with RDP classifications and available metadata were selected (representing 3,551 unique samples; Supplementary Table S1), and 665 samples were prioritized for genome sequencing based on availability, strain novelty compared to existing databases, and body site of origin, with a strategic approach used to avoid sequencing similar samples repeatedly (Supplementary Table S2).

### Genome sequencing, assembly and annotation

DNA extraction and whole genome shotgun sequencing on the Illumina platform was performed as previously described (22). One Button Velvet (Version 1.1.06) (27) was used to assemble the genomes and HMP cut-offs and settings (7) were applied for a sequence to have enough contiguity to be advanced to the annotation level. We screened for contamination by blasting the assembled supercontigs against a database of all HMP strains sequenced in house using BlastN. If supercontigs from the same strain had a top hit against 2 different genera, this strain was considered contaminated and discarded. Those that failed were discarded from the pipeline. The GC% plot was reviewed as an additional measure to detect mixed data from different sources. Any contigs identified as contaminated were removed and the resulting assemblies continued into the annotation pipeline. Annotation was performed as previously described for the HMP reference genomes (22), and taxonomic validation was performed by checking the 16S gene against the reference database, and ensuring that the taxonomic identity of the majority of the predicted genes (by BLASTp against GenBank’s bacterial NR database (31)) matched the identity determined by the RDP.

### Analysis of HMP metagenomics shotgun data

To quantify the value of our 665 newly-assembled HMSC genomes, we mapped 1391 HMP samples from 6 body sites against a baseline bacterial database containing all GenBank complete and drafted bacterial genomes (circa March 2016) (22). We then mapped the same samples to the GenBank database with our 665 HMSC genomes added.

When mapping sample reads to GenBank database entries, we only used GenBank entry genomes with annotation. In cases in which there were multiple entries for a single taxa, we used only the longest representative. The HMP samples used as queries were chosen from 6 body sites: Anterior nares, Buccal mucosa, Posterior fornix, Stool, Tongue dorsum and Supragingival plaque (22). Outliers with regards to sample read counts (‘cleaned’ read counts per sample) were filtered by applying an upper limit of 3x the standard deviation of read counts above the mean per-sample for each body site. A lower cutoff was also set to remove roughly the bottom 5% of the smallest samples. This resulted in 1391 samples being used as queries.

Contaminating human reads in query samples were masked using BMTagger (ftp://ftp.ncbi.nlm.nih.gov/pub/agarwala/bmtagger). Duplicate reads were marked and removed using a modified version of EstimateLibraryComplexity, a tool from the Picard package (http://picard.sourceforge.net/index.shtml). Low quality sequences were then trimmed away using a modified version of the script trimBWAstyle.pl (Fass, J., Unpublished, The Bioinformatics Core at UC Davis Genome Center). This script removed bases with a quality of 2 or less from the ends of reads, an indicator of uncertain quality as defined by Illumina’s End Anchored Max Scoring Segments (EAMMS) filter. After masking and quality trimming, reads with fewer than 60 consecutive non-N bases were removed. The cleaned reads for each sample that passed our filters were then mapped against both databases (GenBank (GB) bacterial genomes & GB bacterial genomes + HMSC genomes) using bowtie2 v2.2.5.

## Supplementary Table Captions

(Tables provided as separate MS Excel files)

**Supplementary Table S1:** Full taxonomy, isolate source, and sequence cluster information for all 3,551 unique 16S samples with RDP classifications and available metadata.

**Supplementary Table S2:** Accession information, full taxonomy, isolate source, and sequence cluster information for all 665 sequenced samples.

## Data Availability

All assembled and annotated genomic sequences and metadata are deposited in NCBI’s Genbank(21). All BioProject, BioSample and Accession IDs are provided per sample in Supplementary Table S2.

## Acknowledgements

This work was supported by grant U54 HG004968.

